# Leptin receptor deficiency suppresses gastric tumorigenesis by limiting stromal activation and tumor microenvironment development

**DOI:** 10.64898/2026.08.21.746140

**Authors:** Kyoko Inagaki-Ohara, Daisuke Motooka, Itaru Yamanaka, Takahiro Nakayama, Shaniya Abudureyimu, Hiroyuki Tezuka, Eiko Sakurai, Kaori Ushida, Takuya Kato, Shizuko Nagao, Yasuhiko Minokoshi, Akihiko Yoshimura, Atsushi Enomoto, Naoya Asai

## Abstract

Leptin receptor (LEPR) signaling has been implicated in multiple malignancies; however, its role in gastric tumors remains poorly defined. We previously demonstrated that mice with gastrointestinal epithelial cell-specific deletion of suppressor of cytokine signaling 3 (SOCS3 cKO), a negative feedback regulator of LEPR signaling, develop gastric tumors due to aberrant leptin production and LEPR activation. Here, we demonstrate that concurrent deletion of both *Socs3* and *Lepr* (double knockout; DKO) under the same promoter substantially suppresses gastric tumorigenesis and markedly prolonged survival. Whereas SOCS3 cKO mice exhibited early stromal activation, increased TGF-β1 production, accumulation of cancer-associated fibroblasts (CAFs) and collagen deposition, these tumor-promoting alterations were substantially attenuated in DKO mice. Additionally, DKO mice showed reduced inflammatory cytokine and chemokine signaling, decreased the accumulation of Gr-1^+^CD11b^+^ myeloid-derived suppressor cells, and reduced LEPR and TGF-β signaling. Analysis of The Cancer Genome Atlas stomach adenocarcinoma cohort revealed high *LEPR* expression in the chromosomal instability and genomically stable subtypes, correlating with poor prognosis. Moreover, *LEPR* expression was mutually exclusive with *CLDN18* and *ERBB2*, two major therapeutic biomarkers, and positively correlated with a CAF-related transcriptional signature. Our findings identify LEPR signaling in epithelial cells as a key driver of gastric tumorigenesis through promotion of stromal activation and tumor microenvironment development. They further highlight LEPR as a promising therapeutic target for patients with gastric cancer who are unlikely to benefit from current ERBB2/HER2- or CLDN18-directed therapies.

## Introduction

Gastric cancer remains a major cause of cancer-related mortality worldwide. More than one million individuals are diagnosed annually, and overall survival remains poor despite advances in surgery, systemic therapy, and the widespread *Helicobacter pylori* eradication. This emphasizes the urgent need to discover new molecular drivers and therapeutic targets beyond the conventional pathways.

Leptin is a well-established regulator of appetite through its actions on hypothalamic neurons [1], but it also promotes cell proliferation, angiogenesis, and tumorigenesis [2, 3]. Leptin receptor (LEPR) signaling is mediated through the JAK/STAT, PI3K/AKT, and ERK pathways and is negatively regulated by suppressor of cytokine signaling 3 (SOCS3). Dysregulated LEPR signaling has been implicated in tumorigenesis across multiple tissues. Pluripotency factors, such as OCT4 and SOX2, directly regulate LEPR expression, and LEPR signaling further reinforces these factors through STAT3 [4]. Consistent with this, pluripotency markers expression is reduced in induced pluripotent stem cells derived from *db/db* mice carrying a mutation in the signaling-competent *Lepr* isoform [5]. In addition to the maintenance of stemness function, LEPR signaling also promotes epithelial-mesenchymal transition (EMT) in gastric cancer cells [6]. Furthermore, LEPR signaling plays a dual role, directly driving tumor cell proliferation and actively suppressing the tumor immune microenvironment [7]. While LEPR signaling crucially drives tumor progression, stemness, and immunosuppression through feedback loops with pluripotency factors, the specific function of locally produced leptin in the gastric mucosa remains largely unexplored.

A fibrotic tumor microenvironment (TME) drives tumor progression across various cancers. Within the extracellular matrix (ECM). Interactions between tumor cells and cancer-associated fibroblasts (CAFs) suppress antitumor immunity, thereby enhancing tumor cell survival, growth [8, 9]. LEPR signaling also contributes to tissue fibrosis in multiple organs [10–12]. In colorectal cancer and cardiac tissue, specific LEPR-expressing cell populations proliferate to promote tumor growth and tissue remodeling, respectively [13, 14]. Chronic tissue injury or inflammation, including *H. pylori* infection or nonsteroidal anti-inflammatory drug exposure, induces dysregulated wound-healing responses, leading to gastric mucosal fibrosis [15, 16]. However, the key molecular mediators and mechanisms underlying initiation of gastric fibrosis remain incompletely understood.

We previously demonstrated that gastrointestinal epithelial cell-specific deletion of *Socs3* promotes gastric tumorigenesis through aberrant leptin production and LEPR signaling in the gastric mucosa [17]. However, because SOCS3 also regulates signaling downstream of multiple IL-6 family cytokines, the specific contribution of LEPR signaling to tumor initiation and progression has remained unclear. To address this question, we generated mice with gastrointestinal epithelial cell-specific deletion of both *Socs3* and *Lepr*. We found that *Lepr* deletion markedly suppresses gastric tumorigenesis, attenuates stromal responses, and reduces the accumulation of CAFs. Furthermore, analysis of The Cancer Genome Atlas (TCGA) data revealed that higher expression of *LEPR* is strongly associated with poor prognosis in patients with gastric cancer. These findings identify LEPR as a key driver of gastric tumorigenesis and a potential therapeutic target.

## Materials and methods

### Mice

B6.129 P2-Lepr^tm1Rck^/J mice (*Lepr*^flox/flox^), which carry homozygous floxed exon 1 *Lepr* alleles (Stock #008327), were purchased from The Jackson Laboratory (ME, USA). To generate mice with gastrointestinal epithelial cell-specific deletion of both *Socs3* and *Lepr* (double-knockout; DKO), T3b-*Socs3* conditional knockout (SOCS3 cKO) mice, generated by crossing T3b-*Cre* transgenic mice with crossing to *Socs3* ^flox/flox^ mice as previously described [17], were crossed with *Lepr*^flox/flox^ mice (The Jackson Laboratory, Stock #008327). The resulting T3b-*Cre Socs3*^flox/+^ *Lepr*^flox/+^ and *Socs3*^flox/+^*Lepr*^flox/+^ mice were intercrossed to generate T3b-*Cre Socs3*^flox/flox^ *Lepr*^flox/flox^ mice and *Socs3* ^flox/flox^ *Lepr*^flox/flox^ littermate controls (Fig. 1A). Polymerase chain reaction (PCR) for *Cre*, *Socs3*, and *Lepr* was performed as previously described [17, 18]. All animal care and experiments were approved by the Animal Ethics Committee of the Prefectural University of Hiroshima and Fujita Health University. All mice were maintained under specific pathogen-free conditions.

**Fig. 1.**
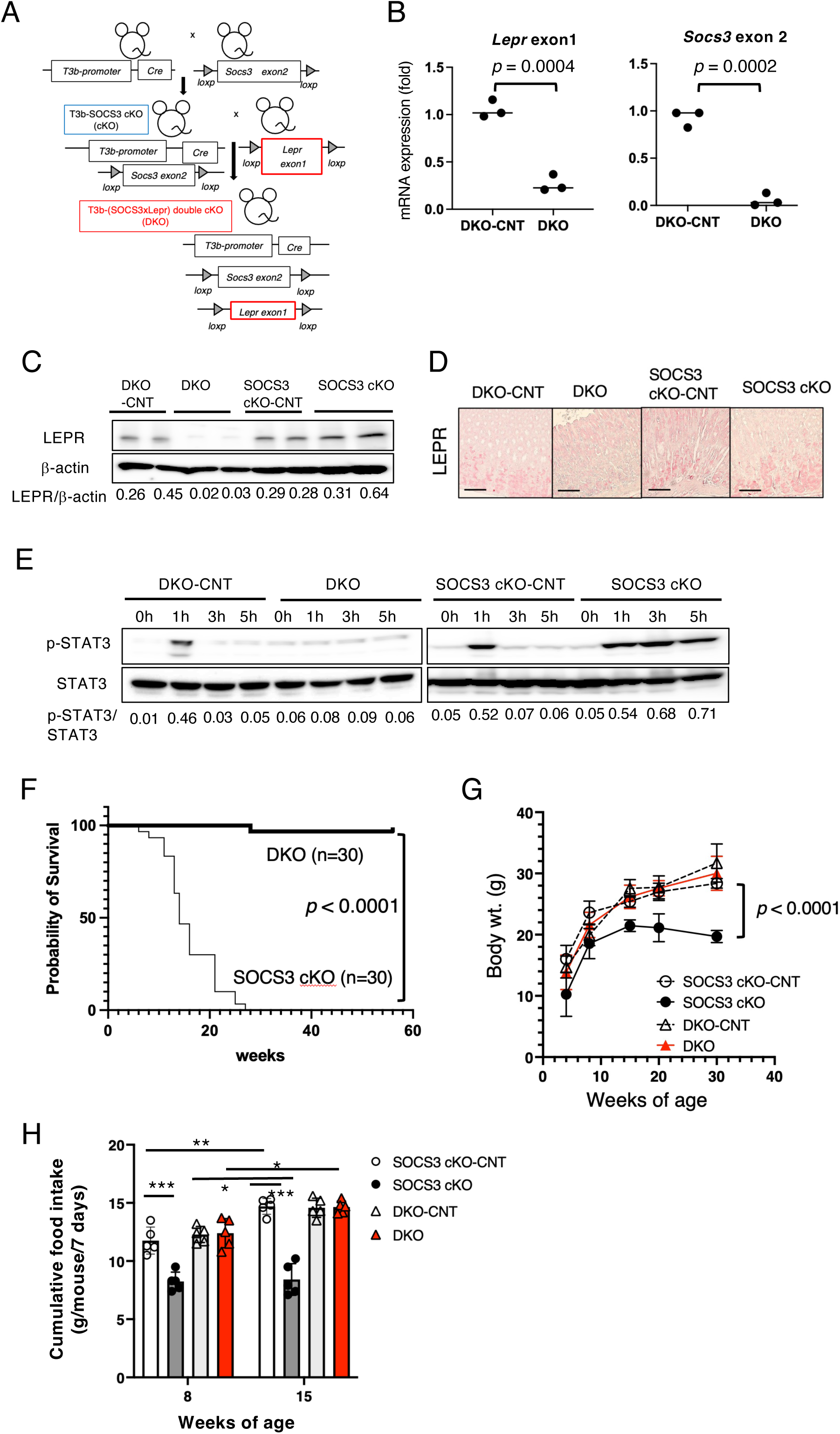
*Lepr* deletion in SOCS3 cKO mice suppresses tumor-associated mortality. (A) Generation of DKO mice. (B) Relative mRNA expression levels of *Lepr* exon 1 and *Socs3* exon 2 in the gastric epithelial cells of DKO and littermate control (DKO-CNT) mice. Statistical significance was determined using an unpaired Student’s *t*-test (n = 3 per group). Individual *p*-values are indicated in the figure. (C) Western blot analysis of LEPR protein expression in the gastric epithelium of DKO, SOCS3 cKO, and their respective littermate controls (DKO-CNT, SOCS3 cKO-CNT). (D) Representative immunohistochemical staining of LEPR in the gastric mucosa. Scale bar = 100 μm. (E) Western blot analysis of phosphorylated and total STAT3 levels in the gastric mucosa of 5-week-old SOCS3 cKO, DKO, and their littermate control mice following intravenous administration of recombinant leptin for the indicated time periods. (F) Kaplan–Meier survival curves of SOCS3 cKO and DKO mice. Overall survival was monitored for up to 58 weeks. Statistical significance was assessed using the log-rank (Mantel-Cox) test. *P*-values is indicated in the figure. (G) Body weight changes for 30 weeks. Statistical analysis was performed using two-way analysis of variance (ANOVA) followed by Sidak’s post-hoc test for multiple comparisons. Statistical significance was assessed using the log-rank (Mantel-Cox) test. *P*-values is indicated in the figure. (H) Food intake at 8 weeks and 15 weeks of age. Statistical analysis was performed using the two-way ANOVA followed by Tukey’s post-hoc test for multiple comparisons. \**p* < 0.05, \*\**p* < 0.001, \*\*\**p* < 0.0001.

### Recombinant leptin administration

To assess leptin-induced STAT3 activation in the gastric mucosa, 5-week-old DKO, SOCS3 cKO and respective control mice were intravenously injected with 5 μg of leptin (FUJIFILM WAKO, Osaka, Japan). Gastric mucosal lysates were prepared at defined intervals for Western blot analysis.

### Preparation of gastric mucosal cells

Gastric mucosal cells were isolated using a modified protocols based on previous studies [19, 20]. Briefly, fresh stomach mucosa was washed with cold PBS containing 1 mM dithiothreitol (DTT), minced into small pieces, and digested with 60 U/mL collagenase (FUJIFILM Wako Pure Chemical, Osaka, Japan). After filtering through a cell strainer to remove cell aggregates and debris, the cell suspension was subjected to discontinuous Percoll (Cytiva, Marborough, MA, USA) density gradient (25%, 40%, and 75%) centrifugation at 600 × *g* for 20 min at room temperature. Epithelial cells were gathered at the 25%–40% interface, while leukocytes were collected at the 40%–75% interface.

### Quantitative PCR (qPCR)

Total RNA was extracted from epithelial cells in gastric epithelial cells using the RNeasy Mini Kit (Qiagen, Hilden, Germany) according to the manufacturer’s instructions. cDNA synthesis was performed using the ReverTraAce® qPCR RT Kit (TOYOBO, Co., Ltd., Osaka, Japan). qPCR was conducted using KOD SYBR qPCR Mix (TOYOBO, Osaka, Japan) with validated primer sets. Primer sequences used for the amplification of *Lepr* exon1, *Socs3* exon 2, and 18S rRNA are as follows: *Lepr* exon 1, forward 5′-GTG TAC ACC TCT GAA GAA AGA TGA-3′ and reverse 5′-CCC AGT GTA ACA AAA CCA CAT AG-3′ [21]; *Socs3* exon 2, forward 5′-CCT CAA GAC CTT CAG CTC CA-3′ and reverse 5′-TGG ATG CGT AGG TTC TTG GT-3′; 18S rRNA forward 5’-AAA CGG CTA CCA CAT CCA AG-3’ and reverse 5’-CCT CCA ATG GAT CCT CGT TA-3’. Amplification was performed on an AriaMx Real-Time PCR System (version 2.0, Agilent Technologies, Foster City, CA, USA). Relative gene expression was calculated using the 2^-ΔΔCt^ method with the 18S rRNA gene used as the internal reference gene.

### Western blot analysis

Protein lysates were prepared from the gastric tissue blot analysis as previously published described [22]. Samples were lysed in lysis buffer containing150 mM NaCl, 0.5% NP-40, and 50 mM Tris-HCl supplemented with protease and phosphatase (cOmplete Mini EDTA-free; Roche Diagnostics GmbH, Mannheim, Germany) inhibitors (PhosStop; Roche Diagnostics GmbH). Lysates were mixed with sodium dodecyl sulfate (SDS) sample buffer, heated at 95 ℃ for 7 min, and sonicated for 1 min. Equal amounts of protein samples were separated by SDS-PAGE and transferred onto polyvinylidene difluoride membranes (Immobilon-P, Merck Millipore, Billerica, MA, USA). The membranes were incubated overnight with primary antibodies against the indicated phosphorylated and total proteins. Following primary antibodies, the membrane was washed with TBST and incubated with a horseradish peroxidase (HRP)-conjugated secondary antibody and were visualized using Chemi-Lumi One detection kit (Nacalai tesque, Kyoko, Japan). The chemiluminescent signals were captured using ImageQuant 800 (Cytiva). The antibodies used for western blotting are summarized in Supplementary Table 1. Band intensities were quantified using ImageJ (version 1.53K).

### Antibodies

Antibodies used for Western blot and immunohistochemical analyses are listed in Supplementary Table 1.

### Immunohistochemical analysis

Tissues fixed in 10% neutral-buffered formalin were paraffin-embedded, sectioned, and stained with hematoxylin and eosin (H&E). Immunohistochemical staining was performed as previously described [17, 22]. Briefly, antigen retrieval was performed using the Retrievagen A kit (BD Biosciences, San Jose, CA, USA), and endogenous peroxidase activity was blocked with 3% hydrogen peroxide in methanol. Sections were incubated overnight at 4°C with primary antibodies (Supplementary Table 1), followed by either biotinylated secondary antibodies and streptavidin–peroxidase or alkaline phosphatase-conjugated secondary antibodies (Histofine SAB-PO kit or Histofine Simple Stain AP; Nichirei Biosciences Inc., Tokyo, Japan). Peroxidase and alkaline phosphatase activities were visualized using ImmPACT DAB and ImmPACT Vector Red, respectively (Vector Laboratories, Burlingame, CA, USA). Sections were counterstained with hematoxylin, and images were acquired using an Axio Imager 2 microscope equipped with an Axiocam 305 camera (Carl Zeiss, Oberkochen, Germany).

### Flow cytometric analysis

Dead cells in the isolated gastric mucosal leukocytes were excluded using 7-aminoactinomycin D (7-AAD) staining, and leukocytes were identified using a Brilliant Violet 510-conjugated anti-mouse CD45.2 monoclonal antibody (clone 104; BioLegend, San Diego, CA, USA). Among the CD45⁺ leukocyte population, Gr-1⁺CD11b⁺ myeloid-derived suppressor cells (MDSCs) and F4/80⁺CD11b⁺ macrophages were identified using a PE-conjugated anti-mouse Gr-1 antibody (clone RB6-8C5; TONBO Biosciences, San Diego, CA, USA), a FITC-conjugated anti-mouse CD11b antibody (clone M1/70; BD Biosciences, San Jose, CA, USA), and an APC-conjugated anti-mouse F4/80 antibody (clone BM8; BioLegend).

### Enzyme-linked immunosorbent assay (ELISA)

Commercial sandwich ELISA kits for gastric leptin and the active form of TGF-β1 were purchased from R&D Systems (Abingdon, OX, UK). Gastric tissue homogenates were prepared according to the manufacture’s protocol provided with the kits. Briefly, gastric tissues were collected, snap-frozen in liquid nitrogen, and stored at −80℃ until use. The tissues were minced into small pieces and homogenized on ice in nine volumes (w/v) of PBS supplemented with a protease inhibitor cocktail (cOmplete Mini EDTA-free; Roche Diagnostics GmbH). The homogenates were centrifuged at 5,000 × g for 15 min at 4℃. The supernatants were immediately collected, aliquoted, and stored at −80℃ until assay. Gastric leptin and the active form of TGF-β1 were measured using sandwich ELISA according to the manufacturer’s instructions.

### Bulk RNA-sequencing (RNA-seq) data analysis

Total RNA was isolated from mouse cells using the RNeasy Mini Kit (Qiagen, Hilden, Germany). Libraries were prepared using the TruSeq Stranded mRNA Sample Prep Kit (Illumina, San Diego, CA, USA) according to the manufacturer’s instructions. Sequencing was performed on an Illumina NovaSeq 6000 platform in 101-base single-end mode. Adapter sequences and low-quality bases were trimmed using Trimmomatic (version 0.38). The processed reads were aligned to the mouse reference genome (mm10) using TopHat (version 2.1.1) in combination with Bowtie2 (version 2.3.5.1) and SAMtools (version 1.11). Raw read counts mapped to each gene were quantified using featureCounts from the Subread (version 2.0.0) with the options -M -O -- fraction. ComBat-ref software [23] was used to perform batch corrections between samples taken at different times during the experiment. Batch-corrected count data were processed for the RLE normalization using DESeq2 software [24]. A pseudocount of 0.125 was added to the normalized count data before a base-2 logarithm was taken. The resulting transformed dataset was used for *t*-test *p*-value and fold-change calculations, as well as for principal component analysis (PCA) and heatmap analysis. PCA plots were generated using the ClustVis web tool [25]. Kyoto Encyclopedia of Genes and Genomes (KEGG) Gene Set Enrichment Analysis (GSEA) was performed using iDEP (version 2.4.0; https://github.com/gexijin/idepGolem/releases/tag/v2.4.0) with a minor modification to the source code to correctly report normalized enrichment scores (NES). Because the heatmaps were generated from log2-transformed expression values, comparisons between the sample_A/sample_B heatmaps were based on differences in log2-transformed values. Heatmaps were generated using R, statistical computing and graphics software (version 4.6.0). The RNA-seq data generated in this study have been deposited in the Gene Expression Omnibus (GEO) database.

### TCGA gastric cancer cohort analysis

RNA-seq and clinical data for the TCGA stomach adenocarcinoma (TCGA-STAD) were obtained from the Genomic Data Commons (GDC) portal (https://portal.gdc.cancer.gov/). The clinical dataset comprised 295 primary gastric adenocarcinomas cases. Clinical annotations were matched to STAR-derived gene expression data using TCGA sample barcodes. Transcripts per million (TPM) expression data were available for 274 primary tumors from the *Nature* 2014 cohort and were transformed as log_2_-transformed as log_2_(TPM + 1). *LEP* and *LEPR* expression levels were compared between the 274 primary tumors and 30 normal gastric tissue samples using the two-sided Wilcoxon rank-sum test. Of the 30 normal samples, 27 were derived from patients who also had a tumor sample in the analyzed cohort. Tumors were stratified according to the Lauren classification (diffuse, intestinal, or mixed) and TCGA molecular subtype (chromosomal instability [CIN], microsatellite instability [MSI], Epstein–Barr virus-positive [EBV], or genomically stable [GS]). Differences among three or more groups were evaluated using the Kruskal–Wallis test. When applicable, pairwise comparisons were performed using two-sided Wilcoxon rank-sum tests with Benjamini–Hochberg correction for multiple comparisons. The expression distribution of *LEPR*, *CLDN18*, and *ERBB2* among the 274 primary tumors was visualized using log_2_(TPM + 1) values. For the extended co-expression analysis, upper-quartile-normalized fragments per kilobase of TPM mapped reads (FPKM-UQ)-derived gene expression Z-scores for 617 cases were downloaded directly from the GDC portal and used without further transformation or standardization. Expression relationships among *LEPR, CLDN18*, and *ERBB2*, between *LEPR* or *CLDN18* and *MYH11, TNC, SYNPO2, SFRP4, ERBB2, DES, ACTG2, and ACTA2*, were visualized using scatter plots. Pearson correlation coefficients and two-sided P-values were calculated, and linear regression lines were fitted. Identical axis limits were applied to all Z-score scatter plots.

### Statistical analysis

Statistical significance between two groups was analyzed using an unpaired Student’s t-test. For multiple-group comparisons or time-course analyses, data were evaluated using one-way or two-way analysis of variance (ANOVA), followed by Tukey’s or Sidak’s post-hoc tests for multiple comparisons. Survival curves were compared using the log-rank (Mantel-Cox) test. All statistical analyses were conducted using GraphPad Prism software (version 10; GraphPad software, San Diego, CA, USA). A *P*-value < 0.05 was considered statistically significant.

## Results

### *Lepr* deletion in SOCS3 cKO mice suppresses gastric tumorigenesis and prevents tumor-associated mortality

To determine whether LEPR signaling directly contributes to gastric tumorigenesis in SOCS3 cKO mice, we generated DKO mice lacking both *Socs3* and *Lepr* in gastrointestinal epithelial cells by crossing SOCS3 cKO mice with *Lepr*^flox/flox^ mice (Fig. 1A). Both mRNA expressions of *Lepr* exon 1 and *Socs3* exon 2 in the gastric epithelial cells were remarkably decreased in DKO mice (Fig. 1B). LEPR protein levels were markedly reduced in DKO mice compared with littermate control (DKO-CNT) mice (Fig. 1C, 1D). Following intravenous leptin administration, SOCS3 cKO mice exhibited sustained STAT3 phosphorylation in the gastric mucosa, whereas this response was markedly attenuated in DKO mice (Fig. 1E). In both littermate control mice (DKO-CNT and SOCS3 cKO-CNT), STAT3 activation was transient. These findings confirm efficient and functional *Cre*-mediated deletion of both *Lepr* and *Socs3* in the gastric epithelium of DKO mice. Ninety-seven % of DKO mice survived beyond 50 weeks of age, whereas all SOCS3 cKO mice died by 30 weeks (Fig. 1F). DKO mice exhibited increased body weight and food intake, whereas SOCS3 cKO mice displayed progressive weight loss beginning at 8 weeks without increased food intake due to gastric tumor development (Fig. 1G, 1H). These results indicate that LEPR signaling is essential for tumor-associated mortality in SOCS3 cKO mice.

### *Lepr* deletion suppresses stromal activation and remodeling during gastric tumor progression

Histopathological analysis revealed that SOCS3 cKO mice developed progressive gastric lesions characterized by early parietal cell loss, stromal expansion, fibroblast proliferation, and mild infiltrating cells with low-grade nuclear atypia by 4 weeks of age (Fig. 2). Between 8 and 15 weeks of age, oxyntic atrophy progressed further and was accompanied by an increased frequency of Ki67-positive proliferating cells, accumulation of Alcian blue-positive acidic mucin, an intestinal-type mucin, and reduced expression of H^+^K^+^-ATPase, a marker of gastric acid-producing parietal cells (Supplementary Fig. S1A). These mice also exhibited advanced fibrosis, disruption of glandular architecture, prominent infiltrating cells, and rapid tumor progression, with some lesions invading the muscle layer by 30 weeks of age (Fig. 2 and Supplementary Fig. S1B). In contrast, DKO mice showed no detectable gastric tumor formation and largely preserved normal gastric architecture even at 30 weeks of age, when all SOCS3 cKO mice had died. Quantitative histopathologic scoring results supported these observations, demonstrating progressive disease severity in SOCS3 cKO mice and marked suppression of gastric lesions in DKO mice (Supplementary Fig. S2). These results indicate that LEPR signaling promotes early stromal activation and subsequent remodeling during gastric tumor progression.

**Fig. 2.**
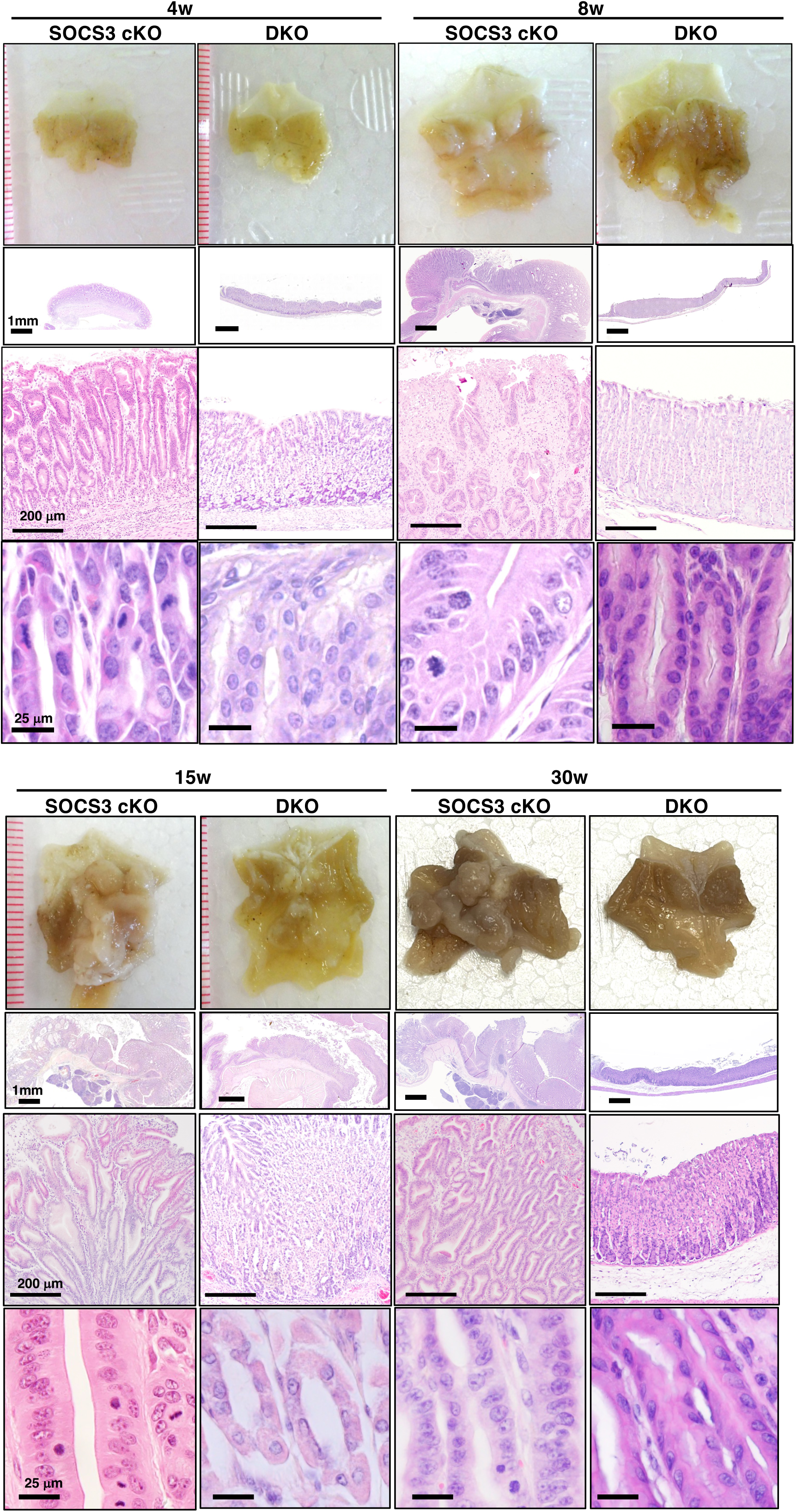
*Lepr* deletion in SOCS3 cKO mice suppresses stromal activation and gastric tumorigenesis. First row: Macroscopic images of the stomachs from SOCS3 cKO and DKO mice at 4, 8, 15, and 30 weeks of age. Second and third rows: Low- and high-magnification images of H&E-stained gastric mucosa. Fourth row: High-magnification images of nuclei corresponding to regions shown in the third row. Representative images from three independent experiments are shown.

### *Lepr* deletion attenuates CAF accumulation and TME formation

Because H&E staining revealed prominent early stromal activation in SOCS3 cKO mice, we examined TME formation in the gastric mucosa. α-SMA–positive fibroblasts were increased in SOCS3 cKO mice at 4 weeks of age, before overt tumor formation, and progressively developed into CAFs by 15 weeks, whereas DKO mice showed no ectopic α-SMA expression (Fig. 3A). Consistent with this, Masson’s trichrome staining revealed extensive collagen deposition in SOCS3 cKO mice but not in DKO mice (Fig. 3B). Hypoxia- and fibrosis-associated molecules, including HIF-1α, TIMP1, and tenascin-C (TNC), were induced at 4 weeks of age and were further elevated at 15 weeks in SOCS3 cKO mice, whereas their expression remained little in DKO mice (Fig. 3C). LEFTY1, a stemness-associated marker [26] in the gastric mucosa, was strongly expressed in SOCS3 cKO mice but faintly detected in DKO mice (Fig. 3D). CD45^+^- immune cell infiltration was observed in SOCS3 cKO mice beginning at 4 weeks and became more prominent between 8 and 15 weeks. In contrast, DKO mice showed minimal immune cell infiltration (Supplementary Fig. S3). Among the infiltrating cells, the absolute number of Gr- 1^+^CD11b^+^ MDSCs were markedly increased in SOCS3 cKO mice by 15 weeks but was substantially reduced in DKO mice (Fig. 4A and Supplementary Fig. S4), whereas the proportion of F4/80^+^CD11b^+^ conventional macrophages differed little between the two groups, although absolute cell number was significantly higher in SOCS3 cKO mice than those in other groups (Fig. 4B).

**Fig. 3.**
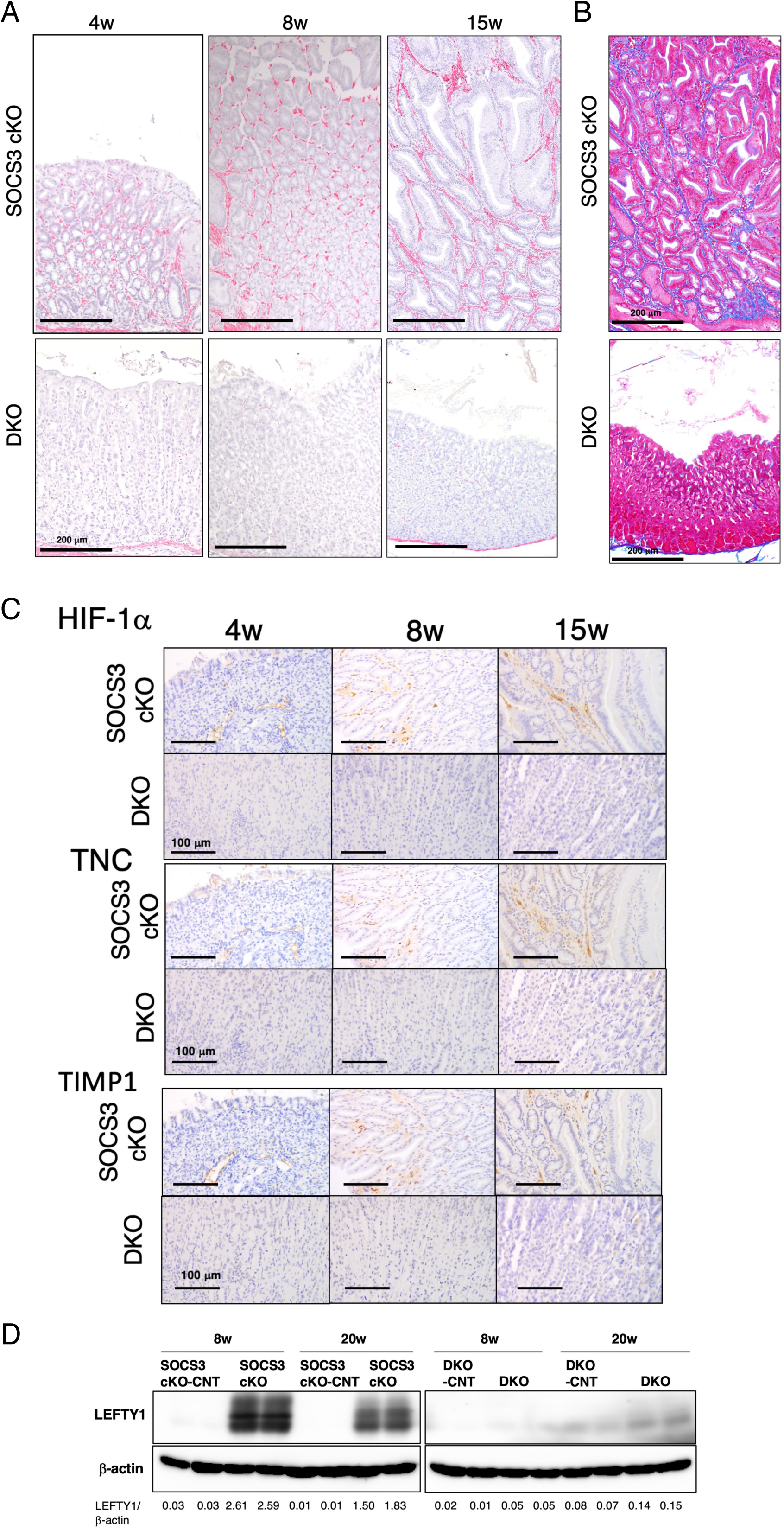
*Lepr* deletion in SOCS3 cKO mice suppresses CAF accumulation and TME development in the gastric mucosa. (A) Representative immunohistochemical staining of α-SMA in the gastric mucosa of SOCS3 cKO and DKO mice at 4, 8, and 15 weeks of age. (B) Representative Masson’s trichrome staining showing collagen deposition in the gastric mucosa of SOCS3 cKO and DKO mice at 15 weeks of age. (C) Representative immunohistochemical staining of HIF-1α, TNC, and TIMP1 in the gastric mucosa of SOCS3 cKO and DKO mice at the indicated ages. Representative images from three independent experiments are shown. (D) Western blot analysis of LEFTY1 expression in the gastric mucosa of SOCS3 cKO, DKO, and their respective littermate controls at 8 and 20 weeks of age. The Western blot signal intensities were quantified using the ImageJ software.

**Fig. 4.**
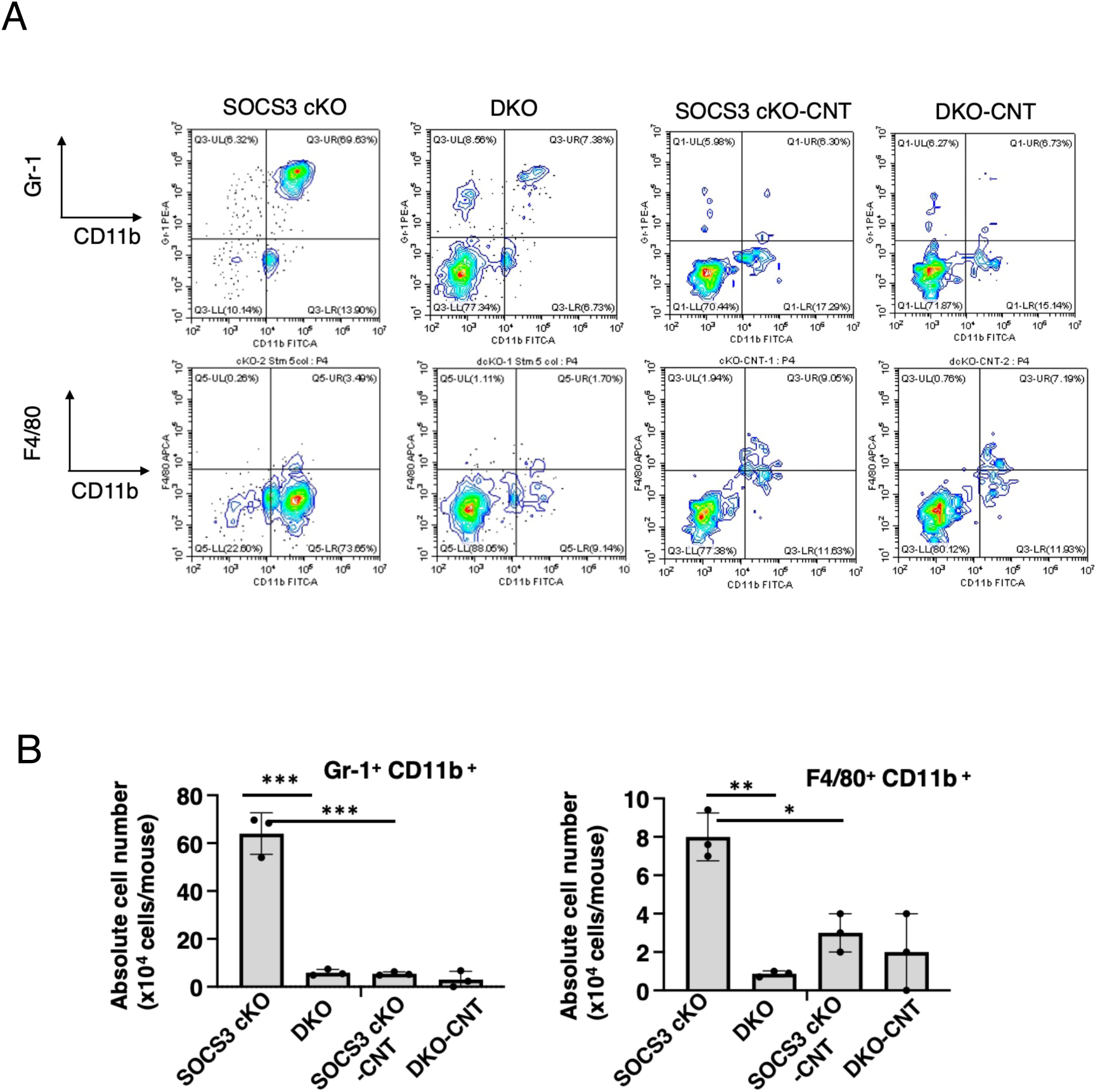
*Lepr* deletion in SOCS3 cKO mice suppresses accumulation of MDSCs in the gastric mucosa. (A) Representative FACS plots showing Gr-1^+^ CD11b^+^ MDSCs and F4/80^+^ CD11b^+^ macrophages in the stomachs of 15-week-old SOCS3 cKO, DKO, and their respective littermate controls. (B) Absolute cell number of MDSCs and macrophages shown in (A). Data are presented as the mean ± SD of three independent experiments, with two mice per group in each experiment. Statistical analysis was performed using one-way ANOVA followed by Tukey’s post-hoc test for multiple comparisons. \**p* < 0.01, \*\**p* < 0.001, \*\*\**p* < 0.0001.

To identify factors associated with CAF accumulation, we next examined TGF-β1 expression, a key profibrotic cytokine, in the gastric tissue. Active form of TGF-β1 in the stomach was detectable at 4 weeks and increased through 15 weeks in SOCS3 cKO mice, whereas DKO mice showed undetectable levels comparable to littermate controls mice (Fig. 5A). Gastric leptin was detected in both the SOCS3 cKO and DKO mice at 4 weeks of age, with a trend toward lower levels in DKO mice (Fig. 5B). In SOCS3 cKO mice, the leptin levels increased until 8 weeks and then tended to decline by 15 weeks, whereas in the DKO mice, its level remained unchanged from 4 to 15 weeks. Consistent with these findings, phosphorylation of SMAD2/3, a major transcription factor of TGF-β signaling, together with phosphorylation of STAT3, AKT, and ERK, key transcription factors of LEPR signaling, was markedly increased in SOCS3 cKO mice but remained substantially lower in DKO mice (Fig. 6). EGFR, which is transactivated by LEPR [22], was highly phosphorylated in SOCS3 cKO mice but faintly detected in DKO mice (Fig. 6). Collectively, these findings indicate that LEPR signaling acts upstream of early TME formation driving TGF-β1 induction, fibroblast activation, hypoxia-associated factors, MDSC accumulation, and enhancement of stemness, thereby contributing to tumor initiation, whereas *Lepr* deletion markedly suppresses these processes.

**Fig. 5.**
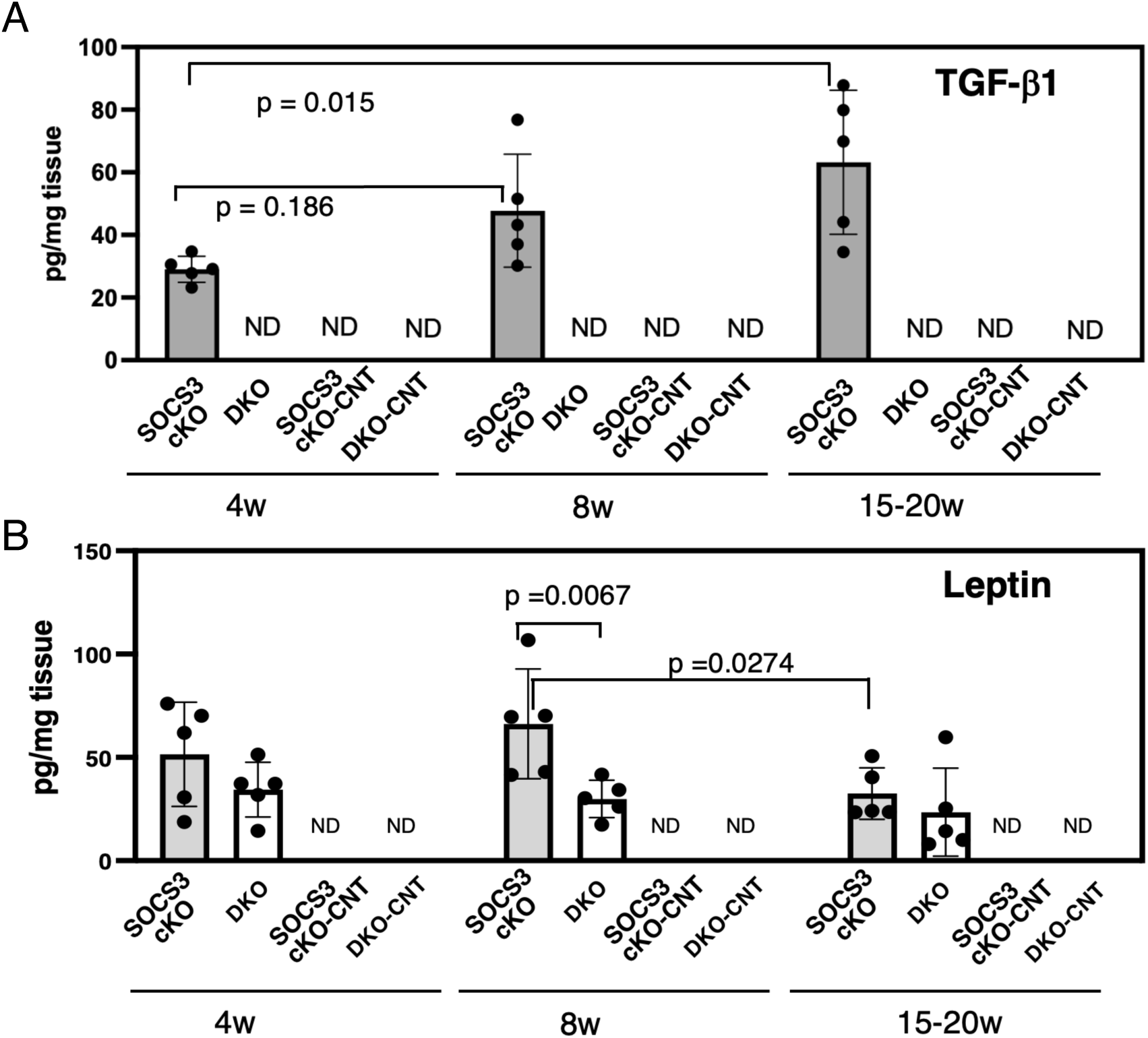
*Lepr* deletion in SOCS3 cKO mice attenuates TGF-β1 production in the gastric tissue. Levels of the active form of (A) TGF- β1 and (B) leptin in the gastric tissue homogenates from SOCS3 cKO, DKO, and respective littermate controls. Data are presented as the mean ± SD (n = 5 mice per group). Active TGF-β1 levels were analyzed using one-way ANOVA followed by Tukey’s post-hoc multiple comparisons, and leptin levels were analyzed using two-way ANOVA followed by Tukey’s post-hoc multiple comparisons. *p*-values are indicated in the figure. ND, not detected.

**Fig. 6.**
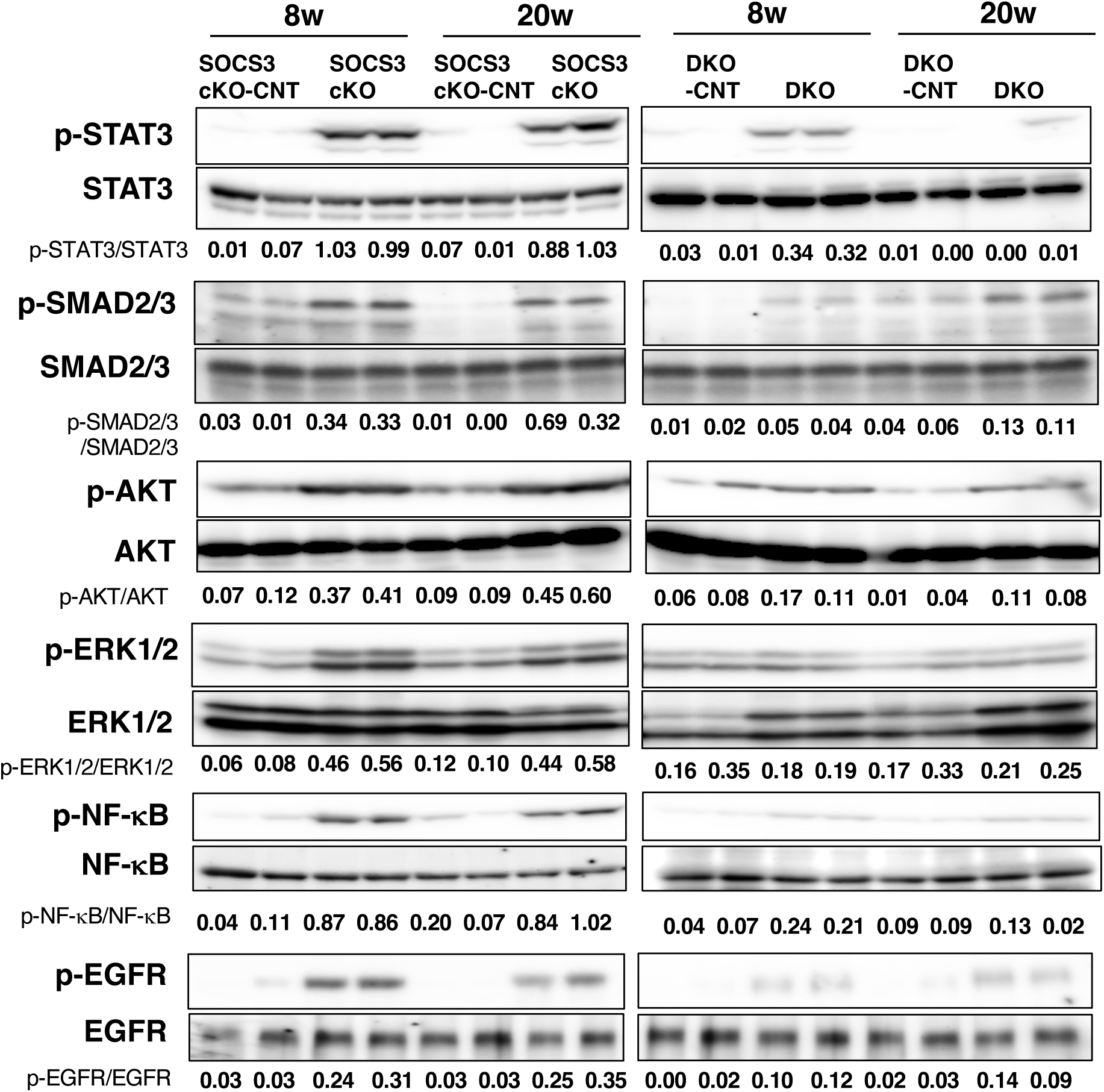
*Lepr* deletion in SOCS3 cKO mice attenuates activation of LEPR- and TGF-β-associated signaling pathways in the gastric mucosa. Western blot analysis and quantification of phosphorylated and total STAT3, SMAD2/3, AKT, ERK1/2, EGFR, and NF-κB expression in the gastric mucosa of 8- and 20-week-old SOCS3 cKO, DKO, and their respective littermate controls. Western blot signal intensities were quantified using the ImageJ software and are represented as the ratio of phosphorylated protein to total protein.

### *Lepr* deletion suppresses inflammatory and stromal reaction pathways during gastric tumorigenesis

Bulk RNA-seq analysis of the gastric mucosa at 8 weeks of age revealed distinct transcriptional profiles among SOCS3 cKO, DKO, and their respective littermate control mice (SOCS3 cKO- CNT and DKO-CNT). PCA showed tight clustering of littermate control groups, whereas SOCS3 cKO and DKO samples clearly separated from the controls and from each other (Fig. 7A). KEGG-based GSEA revealed marked enrichment of cytokine, chemokine, HIF-1, and intracellular signaling pathways in the gastric mucosa of SOCS3 cKO mice (Fig. 7B). Upregulated transcripts included members of the *Il1*, *Il6*, *Il18*, and *Tnf* cytokine families, as well as CXC- and CC chemokine-related genes, including *Cxcl13*, *Cxcr5*, and *Ccl24* (Fig. 7C). Consistent with these findings, phosphorylation of STAT3, NF-κB, AKT, and ERK, key mediators of proinflammatory cytokine and chemokine signaling, was markedly increased in SOCS3 cKO mice but remained substantially lower in DKO mice (Fig. 6). Stromal reaction genes associated with CAF activation, including *Timp1* and *Tnc*, were markedly upregulated in SOCS3 cKO mice but substantially downregulated in DKO mice, consistent with the protein expression (Fig. 3C). In line with the histopathological findings (Fig. 2 and Supplementary Fig. S1A), expression of gastric lineage genes, including *Atp4a* and *Atp4b*, and *Gast*, which encode the H^+^K^+^-ATPase subunits and gastrin, respectively, was markedly reduced in SOCS3 cKO mice (Fig. 7C). In contrast, intestinal-associated genes, including *Sfrp4*, which encodes a key extracellular regulator of Wnt signaling in the intestine, and *Pigr,* which encodes pIgR, responsible for transporting IgA across the epithelial barrier into the intestinal lumen, were highly expressed (Fig. 7C). Taken together, these findings indicate that *Lepr* deletion broadly suppresses inflammatory signaling, stromal activation, and lineage pathways associated with gastric tumorigenesis.

**Fig. 7.**
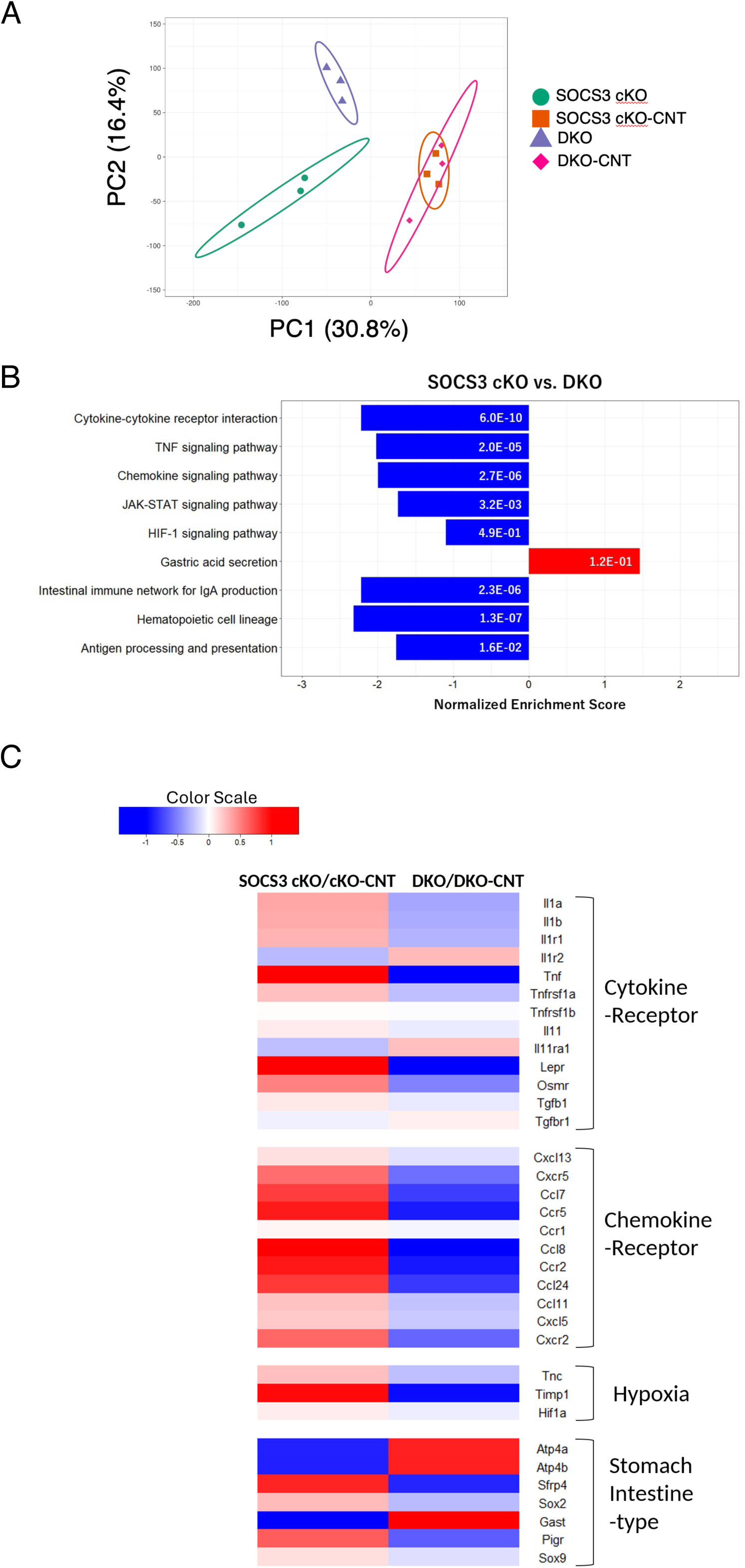
RNA-seq analysis reveals that *Lepr* deletion in SOCS3 cKO mice suppresses inflammatory, stromal activation, and lineage-reprogramming pathways. Gastric tissues were collected from 8- week-old SOCS3 cKO, DKO mice, and their respective littermate control mice (n = 3 mice per group). (A) PCA of bulk RNA-seq data showing distinct clustering of SOCS3 cKO, DKO, and their littermate controls. (B) KEGG-based gene set enrichment analysis (GSEA) using the pre-ranked analysis. Significantly enriched pathways are shown, and adjusted *p*-values are indicated in the figure. (C) Heatmap showing the differential expression of 34 genes across the four experimental groups. These genes were selected from the following four KEGG pathways: Cytokine-Cytokine Receptor Interaction (mmu04060), Chemokine Signaling Pathway (mmu04062), ECM-Receptor Interaction (mmu04512), and HIF-1 Signaling Pathway (mmu04066). Genes from the Gastric Acid Secretion (mmu04971) and Intestinal Immune Networks for IgA Production (mmu04672) were additionally included.

### *LEPR* is highly expressed in clinically relevant human gastric cancer subtypes

To examine the clinical relevance of the mechanism identified in SOCS3 cKO and DKO mice for human gastric cancer, we analyzed GDC-harmonized RNA-seq data from 304 TCGA-STAD case. A subset of tumors exhibited high LEPR expression (Fig. 8A), with higher expression in diffuse-type tumors than in intestinal-type tumors (Fig. 8B). Per the TCGA molecular subtype classification, CIN and GS tumors exhibited elevated *LEPR* expression (Fig. 8C). *ERBB2*, which encodes HER2, and *CLDN18* are among the most important currently established biomarkers for gastric cancer. *LEPR* expression was also detected in gastric cancer samples as well as *ERBB2* and *CLDN18* (Fig. 8D). Notably, LEPR expression was mutually exclusive with both *ERBB2* and *CLDN18* expression (Fig. 8E). Pearson correlation analysis showed that *LEPR* expression was positively correlated with the expression of CAF-associated genes, including *ACTA2*, *ACTG2*, *DES*, *SYNPO2*, *MYH11*, *TNC,* and *SFRP4*, all of which were also upregulated in SOCS3 cKO mice (Fig. 8F). In contrast, CLDN18 expression showed no positive correlation with CAF-associated genes; instead, their high expressions were mutually distinct, with virtually no samples showing concurrent high expression of both. These results indicate that *LEPR* is highly expressed in clinically relevant gastric cancer subtypes, and that LEPR expression is associated with a CAF-related transcriptional signature that parallels the findings in the murine model.

**Fig. 8.**
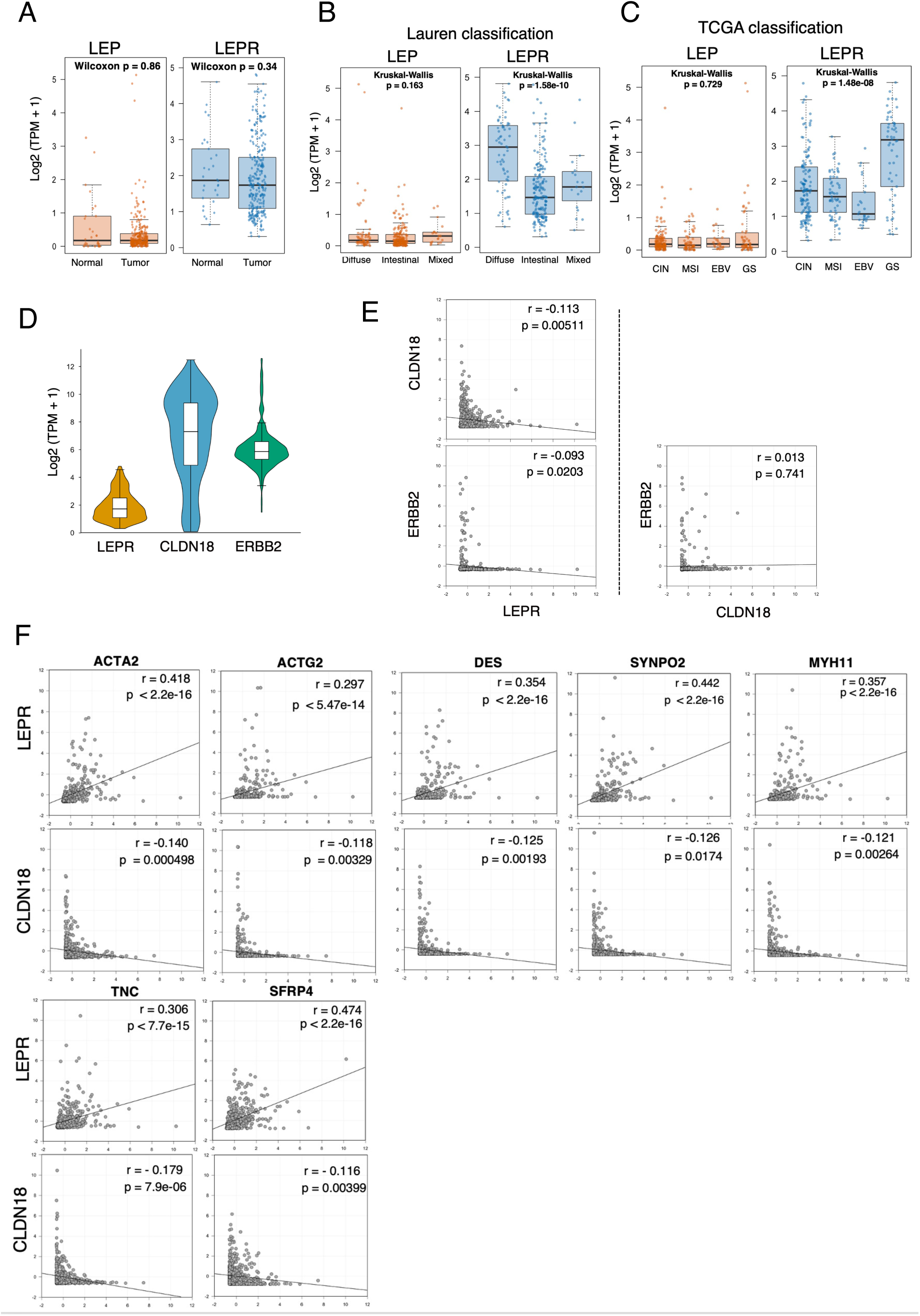
*LEPR* are highly expressed in clinically relevant human gastric cancer subtypes, and *LEPR* expression is associated with a CAF-related transcriptional signature. (A) Comparison of *LEP* and *LEPR* expression levels between primary gastric tumors (n = 274) and normal gastric tissue samples (n = 30) from the TCGA-STAD *Nature* 2014 cohort. Expression levels are shown as log_2_(transcripts per million [TPM] + 1) values. (B) Expression levels of *LEP* and *LEPR* stratified according to Lauren classification: diffuse (n = 64), intestinal (n = 180), and mixed (n = 19). (C) *LEP* and *LEPR* expression levels stratified according to TCGA molecular subtype: chromosomal instability (CIN; n = 136), microsatellite instability (MSI; n = 59), Epstein–Barr virus-positive (EBV; n = 25), and genomically stable (GS; n = 54). (D) Distributions of *LEPR*, *CLDN18*, and *ERBB2* expression in the 274 primary gastric tumors, shown as log_2_(TPM + 1) values. (E) Scatter plots showing expression patterns between *LEPR* and *CLDN18* and between *LEPR* and *ERBB2*, together with the expression relationship between *CLDN18* and *ERBB2*. (F) Scatter plots showing the relationships between *LEPR* (upper panels) or *CLDN18* (lower panels) with CAF- and stromal-associated genes, including *ACTA2*, *ACTG2*, *DES*, *SYNPO2*, *MYH11*, *TNC*, and *SFRP4*. FPKM-UQ-derived gene expression Z-scores used in panels E and F were downloaded directly from the GDC portal and analyzed without further transformation or standardization. Pearson’s correlation coefficients and two-sided P values are shown in each scatter plot.

## Discussion

This study demonstrates that LEPR signaling in the gastric mucosa plays a pivotal and non-redundant role in the initiation and progression of gastric tumorigenesis. By generating mice lacking both *Socs3* and *Lepr* in the gastrointestinal epithelial cells, we provide direct evidence that LEPR signaling is a major mediator of the tumor initiation and promotion induced by SOCS3 deficiency in the gastric mucosa. SOCS3 cKO mice exhibited early stromal activation and TME formation that preceded gastric tumor initiation, whereas these pathological features were substantially suppressed in DKO mice, which showed prolonged survival and preserved body weight and food intake. These findings identify LEPR as a key driver of gastric tumorigenesis in this model. Notably, our findings place LEPR-dependent stromal activation upstream of overt tumor formation, identifying the establishment of a fibrotic microenvironment as an early event in gastric tumorigenesis.

An important conceptual advance of this study is the demonstration that fibrotic stromal remodeling is an early, LEPR-dependent event that precedes overt tumor formation, rather than merely a secondary response to an established tumor. In SOCS3 cKO mice, stromal expansion, accumulation of α-SMA-positive fibroblasts, TGF-β1 production, and induction of the hypoxia- and fibrosis-associated molecules HIF-1α, TIMP1, and TNC, which further contribute to ECM remodeling and tumor progression, were already evident at 4 weeks of age, before the development of overt gastric tumors. These early stromal changes subsequently progressed to extensive collagen deposition, immune-cell infiltration, and disruption of the normal glandular architecture. Importantly, epithelial *Lepr* deletion markedly suppressed these early alterations and prevented subsequent tumor development. Thus, the temporal sequence of these events, together with their genetic dependence on LEPR, supports a model in which epithelial LEPR signaling establishes a pre-tumorigenic, fibrosis-rich stromal niche that facilitates the subsequent initiation and progression of gastric tumors. This finding suggests that CAF activation and extracellular matrix remodeling are not simply consequences of gastric tumor growth but may constitute essential early steps in LEPR-driven tumorigenesis.

In addition to progressing fibrosis, LEPR signaling coordinately orchestrated inflammatory, immunosuppressive, and lineage-reprogramming pathways within the gastric mucosa. RNA-seq analysis revealed that SOCS3 deficiency led to the upregulation of cytokine pathways (*Il1*, *Il6*, *Il18*, and *Tnf*) alongside with broad induction of chemokine genes, including *Cxcl13*, *Cxcr5*, and *Ccl24*. as well as marked upregulation of CAF-associated stromal remodeling genes, including *Timp1* and *Tnc*. Concurrently, CAF-associated stromal remodeling genes, such as *Timp1* and *Tnc*, were markedly upregulated. Consistent with these transcriptomic profiles, SOCS3 cKO mice exhibited CD45^+^ immune cell infiltration, accumulation of Gr-1^+^CD11b^+^ MDSCs, and induction of HIF-1α, TIMP1, and TNC. Strikingly, these pathological alterations were almost completely abolished by the concomitant deletion of *Lepr*. The CXCL13–CXCR5 axis is a well-established regulator of immune cell recruitment and tumor–immune interactions in malignancy [27, 28]. Furthermore, elevated CD11b^+^ myeloid cell infiltration correlates closely with increased tumor size, venous invasion, lymph node metastasis, advanced TNM stage, and poor postoperative prognosis in gastric cancer [29]. In parallel, SOCS3 cKO mice displayed a profound loss of gastric lineage-specific genes accompanied by the upregulation of intestinal-type genes, including *Pigr* and *Sfrp4*, which are molecular hallmarks of intestinal metaplasia and advanced gastric cancer [30, 31]. Crucially, *Lepr* deletion markedly suppressed the phosphorylation of STAT3, SMAD2/3, AKT, and ERK, the key downstream mediators of LEPR and TGF-β signaling. These findings unequivocally demonstrate that epithelial LEPR signaling serves as a central hub coordinating inflammatory, fibrotic, immunosuppressive, and lineage-reprogramming pathways to establish a tumor-promoting gastric microenvironment. Notably, LEPR transactivates EGFR [22], and hyperactivation of EGFR signaling is intimately linked to aggressive progression and poor clinical outcomes in gastric cancer [32]. Characteristically, the robust EGFR activation observed in SOCS3 cKO mice was completely abolished in DKO mice, conclusively establishing LEPR as a therapeutic master switch to simultaneously deactivate both pathogenic signaling networks.

Clinical evidence corroborates that both leptin and LEPR are highly overexpressed in diffuse- and intestinal-type gastric cancers, directly correlating with deeper tumor invasion, advanced TNM staging, and dismal patient prognosis. [33–38]. Our current findings provide a mechanistic explanation for these clinical observations, demonstrating that *Lepr* deletion effectively obliterates fibrosis, inflammatory cytokine/chemokine signaling, MDSC accumulation, and stemness-associated features in the gastric mucosa. Notably, TCGA data analysis revealed that *LEPR* expression is mutually exclusive with *CLDN18* and *ERBB2*/HER2, the two prominent therapeutic biomarkers in gastric cancer. Instead *LEPR* enrichment was highly restricted to the CIN and GS molecular subtypes, which are frequently detected in diffuse-type tumors and manifest significantly poorer overall survival compared to the EBV and MSI subtypes [39, 40]. Furthermore, *LEPR* expression positively correlated with *ACTA2*, *ACTG2*, *MYH11*, *DES*, and *SYNPO2* genes that precisely define myofibroblastic CAFs, a distinct CAF subtype characterized by massive ECM production within the TME and poor clinical outcomes [9, 41]. Taken together, these findings provide compelling evidence that LEPR-high gastric cancers constitute a biologically distinct, stroma-rich subgroup, wherein the leptin–LEPR axis critically drives CAF-associated ECM remodeling and immune evasion, independently of conventional ERBB2- or CLDN18-mediated oncogenic pathways.

### Conclusions

In summary, our integrated analyses leveraging gene-targeted mouse models and human TCGA datasets firmly establish LEPR as a master driver of gastric tumor initiation and progression. Mechanistically, epithelial LEPR signaling functions as a critical upstream instigator that triggers TGF-β1 production, CAF activation, ECM remodeling, inflammatory cytokine/chemokine cascades, MDSC accumulation, and lineage reprogramming, thereby constructing a fibrotic and immunosuppressive microenvironment tailored for gastric tumorigenesis. In human gastric cancer, this LEPR-driven axis is predominantly exploited within the CIN and GS subtypes. Collectively, these findings highlight the leptin–LEPR axis as a biologically distinct pathway and a promising therapeutic target for gastric cancer patients, particularly those currently left with no viable molecularly targeted treatment options.

## Acknowledgments

We would like to thank all of members of the Division of Host Defense (Inagaki-Ohara Lab), Prefectural University of Hiroshima. In particular, Mr. Seiya Arita for establishing the DKO mouse; Mr. Takumi Ogawa and Mr. Masaharu Okazaki for their contributions to the histological analyses; Mr. Yuta Kinoshita for the histological experiments and careful transport of all samples required for this study; and Mr. Tokisada Nakamura, Mr. Yuki Kuriyama, and Ms. Shino Hachiya for their dedicated care of the experimental mice until the very end. We also thank Mr. Kozo Uchiyama (Department of Pathology, Nagoya University) for preparing paraffin sections; Dr. Aya Yoshimura, Dr. Masanori Kugita, Dr. Yuka Kamei, and Ms. Miwa Sakata for introduction and maintenance of mouse colonies; Dr. Keisuke Hitachi (Division for Therapies Against Intractable Diseases) for help maintenance of our mice in Fujita Health University; Professor Hodaka Fujii and Dr. Toshitsugu Fujita (Hirosaki University, School of Medicine) for critical discussions; and the Research Equipment Sharing Center, Nagoya University Graduate School of Medicine, for instrument availability and use of qPCR instrumentation; Prof. Goro Matuzaki (University of the Ryukyus) for reviewing the manuscript and for valuable comments.

## Author contributions

K. I-O. designed and performed the experiments, analyzed and generated data, and wrote and edited the manuscript. D. M., I. Y., and T. N. performed the RNA-seq analysis. I. Y. generated RNA-seq data, wrote the manuscript. D. M. performed TCGA database analyses, generated data, wrote. H. T. set-up condition of the FACS experiment and discussed the manuscript, and S. A. assisted with FACS data analysis. E. S. generated histological data, and K. U., and N. A. discussed to generate histological data, T. K. critically discussed the results. S. N. strongly supported the breeding of mice. Y. M. advised on the information of *Lepr* floxed mice and assisted with colony maintenance. A. Y. critically critical discussed, review and edited. A. E. discussed and supported the research plan and review and edited the manuscript.

## Abbreviations

CAF1: Cancer-associated fibroblast
ECM1: Extracellular matrix
LEPR: Leptin receptor
TME: Tumor microenvironment

## References

1 Zhang Y, Proenca R, Maffei M, Barone M, Leopold L, Friedman JM. Positional cloning of the mouse obese gene and its human homologue. Nature 1994; 372: 425–432.

2 Inagaki-Ohara K. Gastric Leptin and Tumorigenesis: Beyond Obesity. Int J Mol Sci 2019; 20.

3 Kounatidis D, Vallianou NG, Karampela I, Grivakou E, Dalamaga M. The intricate role of adipokines in cancer-related signaling and the tumor microenvironment: Insights for future research. Semin Cancer Biol 2025; 113: 130–150.

4 Feldman DE, Chen C, Punj V, Tsukamoto H, Machida K. Pluripotency factor-mediated expression of the leptin receptor (OB-R) links obesity to oncogenesis through tumor-initiating stem cells. Proc Natl Acad Sci U S A 2012; 109: 829–834.

5 Gupta MK, Vethe H, Softic S, Rao TN, Wagh V, Shirakawa J et al. Leptin Receptor Signaling Regulates Protein Synthesis Pathways and Neuronal Differentiation in Pluripotent Stem Cells. Stem Cell Reports 2020; 15: 1067–1079.

6 Park KB, Kim EY, Chin H, Yoon DJ, Jun KH. Leptin stimulates migration and invasion and maintains cancer stem-like properties in gastric cancer cells. Oncol Rep 2022; 48: 162.

7 Deka G, Park PH. Leptin Signaling at the Crossroads of Obesity, Immunity, and Cancer: Mechanistic Insights and Therapeutic Implications. Biomol Ther (Seoul) 2026; 34: 780– 799.

8 Contreras-Panta EW, Choi E, Goldenring JR. The Fibroblast Landscape in Stomach Carcinogenesis. Cell Mol Gastroenterol Hepatol 2024; 17: 671–678.

9 Qi J, Sun H, Zhang Y, Wang Z, Xun Z, Li Z et al. Single-cell and spatial analysis reveal interaction of FAP(+) fibroblasts and SPP1(+) macrophages in colorectal cancer. Nat Commun 2022; 13: 1742.

10 Petrescu AD, Grant S, Williams E, An SY, Seth N, Shell M et al. Leptin Enhances Hepatic Fibrosis and Inflammation in a Mouse Model of Cholestasis. Am J Pathol 2022; 192: 484–502.

11 Mao Y, Zhao K, Li P, Sheng Y. The emerging role of leptin in obesity-associated cardiac fibrosis: evidence and mechanism. Mol Cell Biochem 2023; 478: 991–1011.

12 Sun XN, Chen S, Zhao S, Funcke JB, Virostek M, Pedersen L et al. Leptin as a key driver for organ fibrogenesis. Sci Adv 2025; 11: eady7904.

13 Larcher V, Fischer A, Zhang L, Almolla O, Chiesa M, Andriani F et al. Leptin Receptor Fibroblasts Are Preferential Contributors to Cardiac Fibrosis. Circ Res 2026; 138: e327701.

14 Kobayashi H, Gieniec KA, Lannagan TRM, Wang T, Asai N, Mizutani Y et al. The Origin and Contribution of Cancer-Associated Fibroblasts in Colorectal Carcinogenesis. Gastroenterology 2022; 162: 890–906.

15 Genta RM. Helicobacter pylori, inflammation, mucosal damage, and apoptosis: pathogenesis and definition of gastric atrophy. Gastroenterology 1997; 113: S51–55.

16 Scarpignato C. Nonsteroidal anti-inflammatory drugs: how do they damage gastroduodenal mucosa? Dig Dis 1995; 13 Suppl 1: 9–39.

17 Inagaki-Ohara K, Mayuzumi H, Kato S, Minokoshi Y, Otsubo T, Kawamura YI et al. Enhancement of leptin receptor signaling by SOCS3 deficiency induces development of gastric tumors in mice. Oncogene 2014; 33: 74–84.

18 Arita S, Inagaki-Ohara K. High-fat-diet-induced modulations of leptin signaling and gastric microbiota drive precancerous lesions in the stomach. Nutrition 2019; 67-68: 110556.

19 Inagaki-Ohara K, Chinen T, Matsuzaki G, Sasaki A, Sakamoto Y, Hiromatsu K et al. Mucosal T cells bearing TCRgammadelta play a protective role in intestinal inflammation. J Immunol 2004; 173: 1390–1398.

20 Suzuki K, Sentani K, Tanaka H, Yano T, Suzuki K, Oshima M et al. Deficiency of Stomach-Type Claudin-18 in Mice Induces Gastric Tumor Formation Independent of H pylori Infection. Cell Mol Gastroenterol Hepatol 2019; 8: 119–142.

21 Takahashi K, Yamada T, Hosaka S, Kaneko K, Asai Y, Munakata Y et al. Inter-organ insulin-leptin signal crosstalk from the liver enhances survival during food shortages. Cell Rep 2023; 42: 112415.

22 Kinoshita Y, Arita S, Ogawa T, Takenouchi A, Inagaki-Ohara K. Augmented leptin-induced trefoil factor 3 expression and epidermal growth factor receptor transactivation differentially influences neoplasia progression in the stomach and colorectum of dietary fat-induced obese mice. Arch Biochem Biophys 2022; 729: 109379.

23 Zhang X. Highly effective batch effect correction method for RNA-seq count data. Comput Struct Biotechnol J 2025; 27: 58–64.

24 Love MI, Huber W, Anders S. Moderated estimation of fold change and dispersion for RNA-seq data with DESeq2. Genome Biol 2014; 15: 550.

25 Metsalu T, Vilo J. ClustVis: a web tool for visualizing clustering of multivariate data using Principal Component Analysis and heatmap. Nucleic Acids Res 2015; 43: W566– 570.

26 Tsubosaka A, Komura D, Kakiuchi M, Katoh H, Onoyama T, Yamamoto A et al. Stomach encyclopedia: Combined single-cell and spatial transcriptomics reveal cell diversity and homeostatic regulation of human stomach. Cell Rep 2023; 42: 113236.

27 Hussain M, Adah D, Tariq M, Lu Y, Zhang J, Liu J. CXCL13/CXCR5 signaling axis in cancer. Life Sci 2019; 227: 175–186.

28 Xu S, Li D, Ning T, Lu Y, Sun Y, Bai H et al. High expression of CXCL13 predicts a favorable response to immunotherapy by upregulating CXCR5+CD8+ T-cell infiltration in gastric cancer. Front Immunol 2025; 16: 1551259.

29 Okita Y, Tanaka H, Ohira M, Muguruma K, Kubo N, Watanabe M et al. Role of tumor-infiltrating CD11b+ antigen-presenting cells in the progression of gastric cancer. J Surg Res 2014; 186: 192–200.

30 Fristedt R, Gaber A, Hedner C, Nodin B, Uhlen M, Eberhard J et al. Expression and prognostic significance of the polymeric immunoglobulin receptor in esophageal and gastric adenocarcinoma. J Transl Med 2014; 12: 83.

31 Li H, Zhao J, Sun J, Tian C, Jiang Q, Ding C et al. Demethylation of the SFRP4 Promoter Drives Gastric Cancer Progression via the Wnt Pathway. Mol Cancer Res 2021; 19: 1454–1464.

32 Ding B, Wan Y, Wu Y, Zhang Z, Ma Y, Wang Z et al. Phosphorylated-EGFR and MMP7 upregulation in gastric cancer: Association with metastasis and poor prognosis. Oncol Lett 2026; 31: 107.

33 Azuma T, Suto H, Ito Y, Ohtani M, Dojo M, Kuriyama M et al. Gastric leptin and Helicobacter pylori infection. Gut 2001; 49: 324–329.

34 Ishikawa M, Kitayama J, Nagawa H. Expression pattern of leptin and leptin receptor (OB- R) in human gastric cancer. World J Gastroenterol 2006; 12: 5517–5522.

35 Zhao X, Huang K, Zhu Z, Chen S, Hu R. Correlation between expression of leptin and clinicopathological features and prognosis in patients with gastric cancer. J Gastroenterol Hepatol 2007; 22: 1317–1321.

36 Geng Y, Wang J, Wang R, Wang K, Xu Y, Song G et al. Leptin and HER-2 are associated with gastric cancer progression and prognosis of patients. Biomed Pharmacother 2012; 66: 419–424.

37 Zhao L, Shen ZX, Luo HS, Shen L. Possible involvement of leptin and leptin receptor in developing gastric adenocarcinoma. World J Gastroenterol 2005; 11: 7666–7670.

38 Lee KN, Choi HS, Yang SY, Park HK, Lee YY, Lee OY et al. The role of leptin in gastric cancer: clinicopathologic features and molecular mechanisms. Biochem Biophys Res Commun 2014; 446: 822–829.

39 Cancer Genome Atlas Research N Comprehensive molecular characterization of gastric adenocarcinoma. Nature 2014; 513: 202–209.

40 Sohn BH, Hwang JE, Jang HJ, Lee HS, Oh SC, Shim JJ et al. Clinical Significance of Four Molecular Subtypes of Gastric Cancer Identified by The Cancer Genome Atlas Project. Clin Cancer Res 2017; 23: 4441–4449.

41 Hashmi SK, Ceron RH, Heuckeroth RO. Visceral myopathy: clinical syndromes, genetics, pathophysiology, and fall of the cytoskeleton. Am J Physiol Gastrointest Liver Physiol 2021; 320: G919–G935.

