## Supplementary material for "Leptin receptor deficiency suppresses gastric tumorigenesis by limiting stromal activation and tumor microenvironment development": Text (Supplementary)

### Supplementary information

#### Materials and Methods

#### Immunohistochemical analysis

Paraffin-embedded sections of tissues fixed in 10% neutral-buffered formalin were stained with periodic acid-Schiff (PAS) and Alcian blue (pH 2.5) or subjected to immunohistochemical staining for Ki67 and $H^+K^+$-ATPase, as described in our previous study [17, 22]. For immunofluorescence staining, the tissue sections incubated with the primary antibodies were subsequently stained with Alexa Fluor 488-conjugated anti-rat IgG antibody and Alexa Fluor 594-conjugated anti-rabbit IgG antibody (ThermoFisher Scientific, Eugene, OR, USA). The sections were then mounted using Permafluor Aqueous Mounting Medium (Thermo Fisher Scientific, MI, USA) containing 4′,6-diamidino-2-phenylindole (DAPI) (Life Technologies, Carlsbad, CA, USA) and visualized using an Axio Imager 2 microscope equipped with an Axiocam 305 camera (Carl-Zeiss).

#### Histopathologic analysis

Stomach sections from the SOCS3 cKO, DKO, and their respective littermate controls were stained with H&E or PAS-Alcian blue. Histopathological progression in the gastric tissue was evaluated using a semi-quantitative scoring system developed to reflect the chronological phenotypic changes observed in the mice (n = 6–10 per group). The scoring criteria were as follows: (1) Cell infiltration: scored on a scale of 0 to 4, where 0 = no substantial alteration; 1 = mild, and localized inflammatory cell infiltration, 2 = moderate infiltration extending into the lamina propria; 3 = marked infiltration throughout the mucosa; and 4 = severe mucosal infiltration with prominent lymphoid follicle formation. (2) Hyperplasia: scored on a scale of 0 to 3, where 0 = no substantial alteration; 1 = mild epithelial proliferation; 2 = moderate with distinct foveolar hyperplasia; and 3 = marked epithelial thickening. (3) Intestinal metaplasia: evaluated using Alcian blue staining to detect acid mucin-positive intestinal metaplasia (IM), an early feature of mucosal transformation. Lesions were scored on a scale of 0 to 3, where 0 = no substantial alteration; 1 = involvement of <20% of the mucosa; 2 = involvement of 20–50% of the mucosa; and 3 = involvement of >50% of the mucosa. (4) Cellular dysplasia: scored on a scale of 0, 6, 7, and 8, where 0 = no substantial alteration; 6 = mild cellular atypia; 7 = moderate cellular atypia with distinct nuclear enlargement; and 8 = severe cellular atypia characterized by marked nuclear pleomorphism and loss of polarity. (5) Architectural dysplasia: scored on a scale of 0, 6, 7, and 8, where 0 = no substantial alteration; 6 = mild glandular crowding; 7 = moderate glandular branching with architectural distortion; and 8 = severe architectural distortion characterized by back-to-back glandular crowding. (6) Tumor staging: scored on a scale of 10, 12, and 13 based on intramucosal tumor burden and the degree of subsequent luminal obstruction: A score of 10 indicated an early intramucosal tumor with localized neoplastic lesions confined to the mucosa; a score of 12 indicated an advanced intramucosal tumor with extensive horizontal or exophytic growth and marked cellular atypia confined to the mucosa; and a score of 13 indicated a terminal obstructive tumor with a large intramucosal tumor mass causing severe gastric luminal obstruction and high mortality before deep invasion.

### Supplementary Figure Legends

**Fig. S1.** *Lepr* deletion in SOCS3 cKO mice suppresses intestinal characteristic and tumor invasion to muscle layer. (A) Representative immunohistochemical staining of Ki67 (upper panels), $H^+K^+$-ATPase (middle panels), and PAS-Alcian blue staining (lower panels), in the gastric mucosa of 8-week-old SOCS3 cKO and DKO mice. (B) Representative H&E-stained section showing invasion in the muscle layer in 30-week-old SOCS3 cKO mice.

**Fig. S2. Histopathologic scoring of gastric lesions in SOCS3 cKO and DKO mice.** Histopathologic scores of the gastric mucosa from SOCS3 cKO, DKO, and their respective littermate controls at 4, 8–15, and 16–30 weeks of age were determined according to the histopathological criteria described in the Supplementary Materials and Methods. Individual dots represent individual mice (n = 6–10 mice per group), and the horizontal bars indicate the mean score for each group. Individual scores were summed to obtain the cumulative histopathologic score.

**Fig. S3. *Lepr* deletion in SOCS3 cKO mice suppresses leukocyte infiltration in the gastric mucosa.** Representative immunohistochemical staining of $CD45^+$ leukocytes (green) in the gastric mucosa of SOCS3 cKO and DKO mice at 4, 8, and 15 weeks of age. Nuclei are counterstained with DAPI (blue). Representative images from at least three independent experiments are shown.

**Fig. S4. *Lepr* deletion suppresses accumulation of MDSCs in the gastric mucosa of SOCS3 cKO mice.** Representative immunohistochemical staining of Gr-1 and CD11b in the gastric mucosa of SOCS3 cKO and DKO mice at 15 weeks of age. Representative images from at least three independent experiments are shown.
