## Supplementary Table 1 for "Leptin receptor deficiency suppresses gastric tumorigenesis by limiting stromal activation and tumor microenvironment development"

**Supplemental Table S1. Antibodies used for Western blotting and IHC**

| Name of Antibody | Manufacturer catalog No. |
|---|---|
| p-Akt | Cell Signaling Technology 4060 (Ser473) |
| Akt | Cell Signaling Technology（4691S) |
| ATPase (H+K+) | Abcam (2866) |
| β-actin | Proteintech (66009-1-Ig) |
| CD11b | Abcam (ab133357) |
| CD45 | BD Bioscience (550539) |
| p-EGFR (Tyr1068) | Cell Signaling Technology (3777) |
| EGFR | Cell Signaling Technology (4267) |
| Hif-1α | Santa cruz (sc-13515) |
| Ki67 | ThemoFisher Science 14-5698-82 |
| Lefty | Santa cruz (sc-365845) |
| LEPR | Proteintech (20966-1-AP) |
| Leptin | Abcam (ab16227) |
| Ly-6G(Gr-1) | Santa Cruz (sc-53515) |
| p-NF-kb (p65) | Cell Signaling Technology (3037S) |
| NF-kb (p65) | Santa Cruz (sc-8008) |
| p-p44/p42 MAPK (ERK1/2) | Cell Signaling Technology (4370) |
| MAPK (ERK1/2) | Cell Signaling Technology (4695) |
| Phospho-PI3K p85(Tyr458)/p55(Tyr199) | CST #4228 |
| PI3K p85 | CST #4257 |
| α-SMA | Dako (M0851) |
| p-Smad2/3 | Cell Signaling Technology (#8828) |
| Smad2/3 | Santa Cruz (sc-133098) |
| p-STAT3 | Cell Signaling Technology 9145S |
| STAT3 | Cell Signaling Technology #9139 |
| Tenascin-C | Santa Cruz (sc-25328) |
| TIMP-1 | Santa Cruz (sc-21734) |
