## Supplementary figures and images for "Leptin receptor deficiency suppresses gastric tumorigenesis by limiting stromal activation and tumor microenvironment development"

### Supplementary Figures

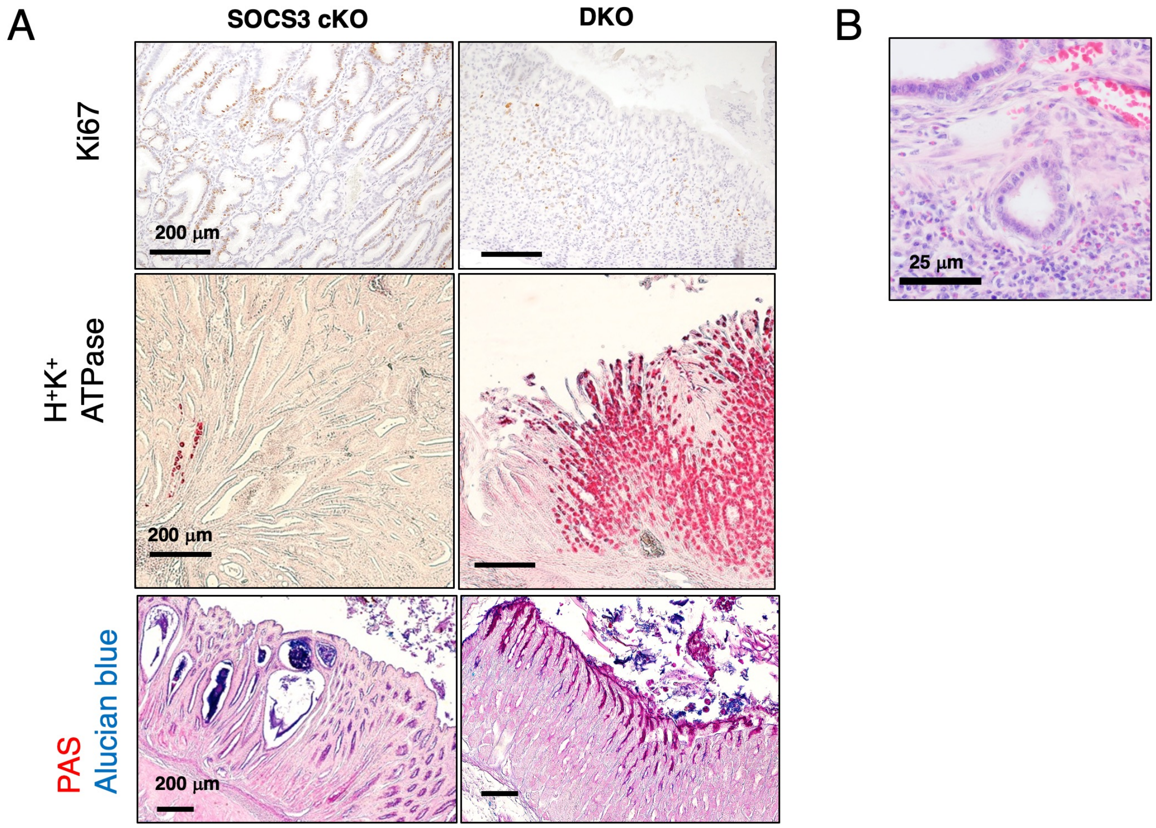


Supplemental Fig. S1

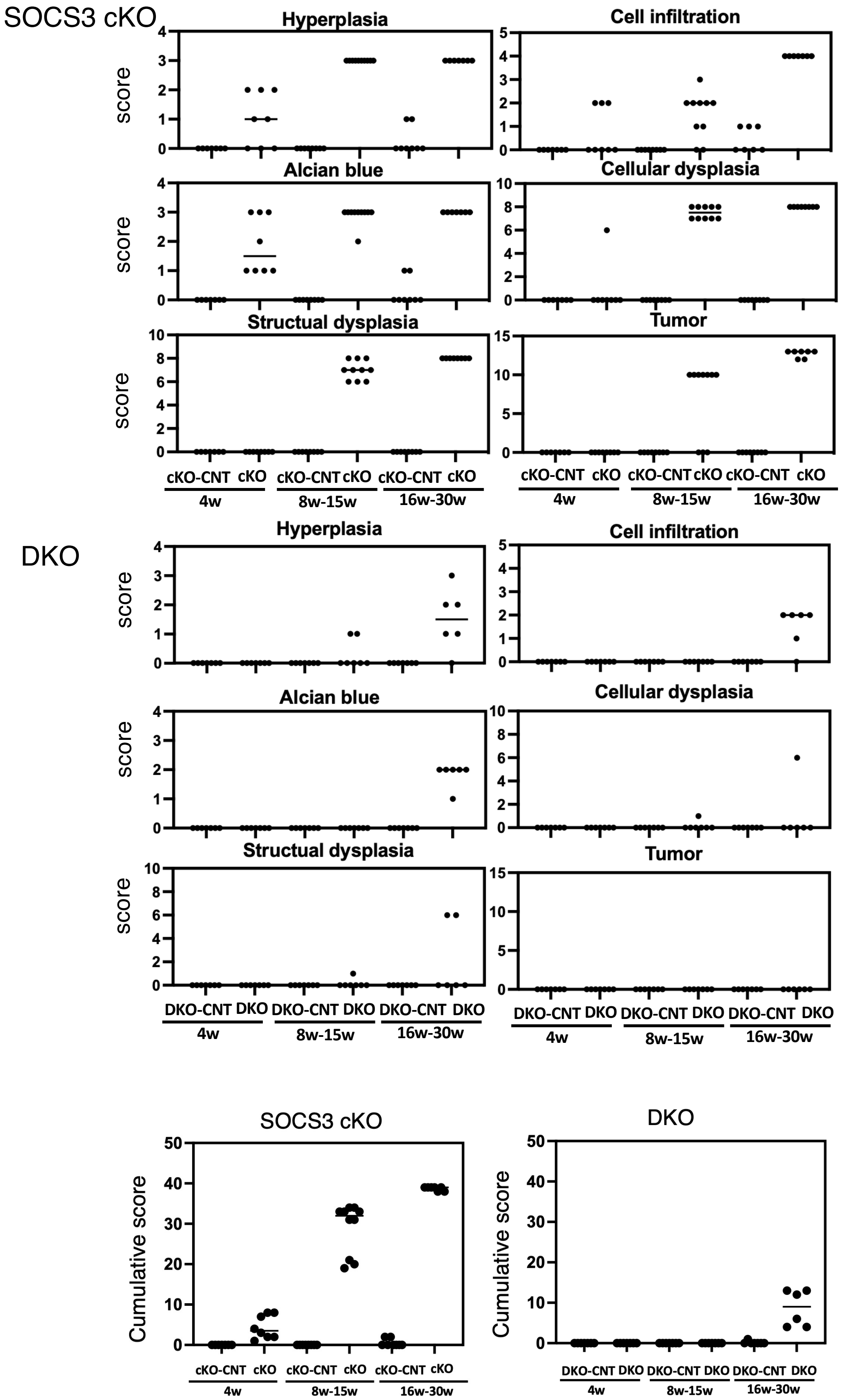


Supplemental Fig. S2

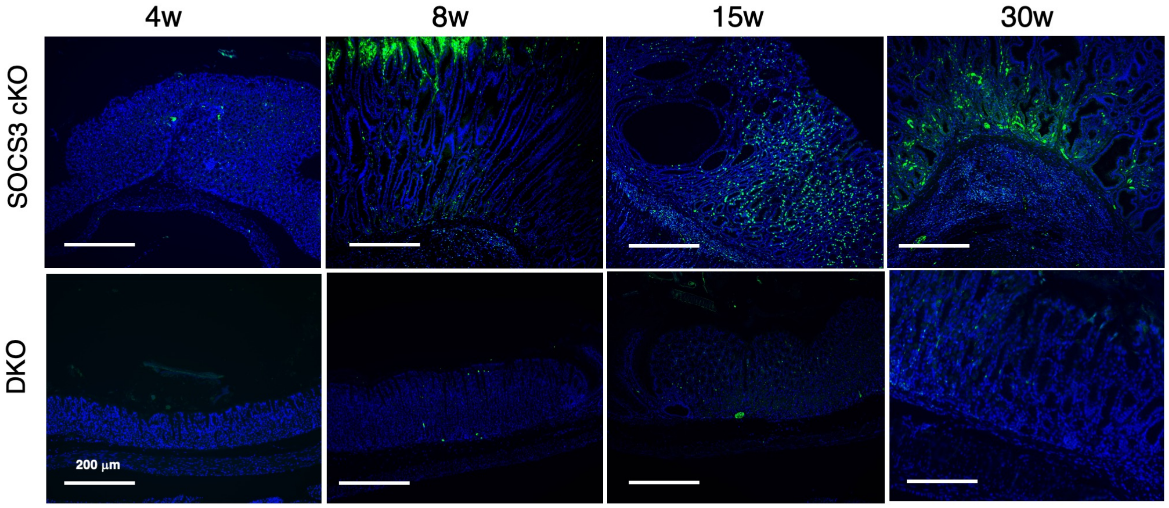


Supplementary Fig. S3

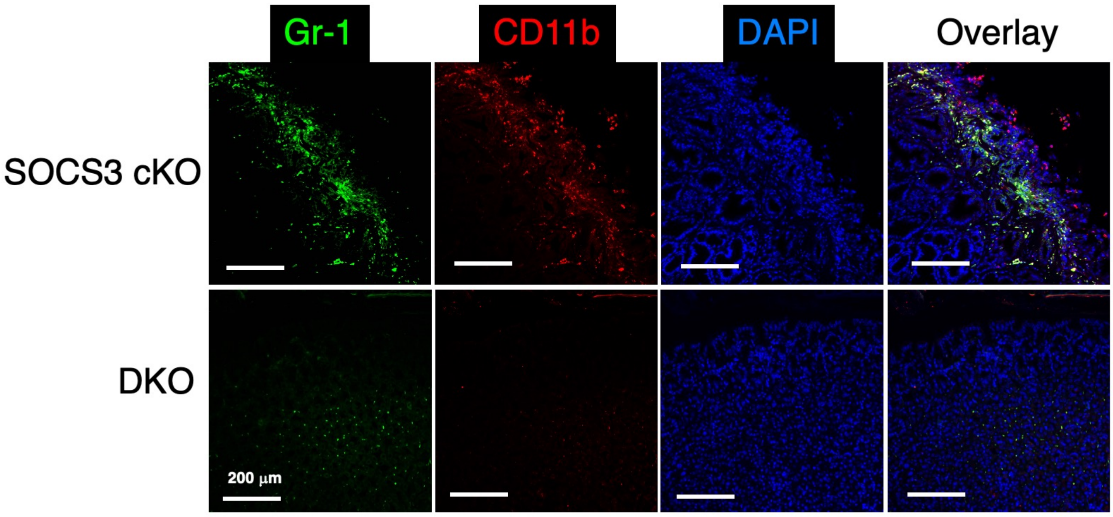


Supplemental Fig. S4
